# Ocean depth modifies host-microbiome ecology and evolution

**DOI:** 10.64898/2026.10.22.761184

**Authors:** Cristina Díez-Vives, Marta Turon, Miguel Lurgi, Lucía Pita, Shaun Nielsen, Almudena Álvarez Fernández, Torsten Thomas, Ana Riesgo, José M Montoya

## Abstract

A major unsolved question in ecology and evolution is how similar free-living and symbiotic microbial communities are. In marine systems, depth homogenises free-living communities due to stronger dispersal and weaker environmental filtering compared to surface waters. We show that symbiotic communities of marine sponges follow a different pattern, with a subset of sponge species mirroring the increased similarity with depth seen in their free-living counterparts, while others remain highly host-specific both in shallow and deep waters. Moreover, in deep waters, more phylogenetically similar hosts harbor more similar microbiomes, yet host-symbiont cophylogeny nearly vanishes. This reveals two contrasting eco-evolutionary assembly rules: in more constant environments like the deep sea, community assembly is driven by host and environmental filtering with minimal signature of coevolution, whereas in variable environments, like shallow waters, host-symbiont coevolution is strong and key to maintain host-specific microbiomes.

## Introduction

Dispersal of individuals tends to homogenize ecological communities. In the case of microbes, prevailing wisdom states that “*everything is everywhere*, but, *the environment selects*” due to weak dispersal limitation (*1–3*). In the ocean, global circulation of water masses favours microbial dispersal (*4*), and in the absence of environmental heterogeneity, the expectation would be a uniform microbial community across scales. However, both dispersal and environmental context differ markedly between surface and deep waters. Dispersal distances and spatial connectivity are greater in the deep ocean than in shallow waters (*5*, *6*). In addition, the surface ocean is characterised by strong spatial heterogeneity which exerts higher selection on microbial communities, while deep waters are comparatively more homogeneous (*7*, *8*). For free-living microorganisms, the combination of lower spatial connectivity and stronger selection by environmental factors in surface waters promotes greater niche differentiation, specialization, and community turnover than in deep waters, where communities are more similar (*6*, *9–13*).

A major unknown is whether host-associated microbiomes follow the same patterns as free-living microbial communities. If they do, we would expect host-associated microbial communities inhabiting shallow waters to be more specialized and compositionally different across hosts than their deep-water counterparts. Unlike free-living microorganisms, host-associated symbiotic microbes experience a relatively stable environment within their hosts, largely buffered from fluctuations in the surrounding waters or sediments, while also benefiting from greater resource availability (*14*, *15*). In turn, symbiotic microbes can expand their host’s physiological capabilities by enabling access to novel nutrient sources, increasing tolerance to environmental stressors, and protecting against pathogens (*16*). Symbiosis is therefore a major driver of evolutionary novelty and ecological diversity (*17*), often involving reciprocal selection dynamics and coevolutionary processes (*18–20*) that can result in dynamics and interaction patterns markedly different from those observed in free-living microbial communities.

Sponges (phylum Porifera) are an evolutionarily and ecologically relevant model for studying host-microbe interactions in early-diverging metazoans, given that their associations with microbial communities likely originated approximately 890 million years ago (*21–23*). Sponges occur worldwide in both marine and freshwater habitats, ranging from the intertidal zone to hadal depths (*24*). Moreover, sponges display a wide range of strategies with respect to their symbiotic community (*25*, *26*). While some species harbor dense and diverse microbial consortia within their tissues and are thus classified as high microbial abundance (HMA) sponges, others harbor less dense communities and are referred to as low microbial abundance (LMA) sponges. Microbial communities of LMA sponges tend to be more uneven, with a few dominant taxa and a larger proportion of low-abundance members. From a functional perspective, LMA sponges exhibit high pumping rates and rely primarily on their own metabolism for nutrition, whereas HMA sponges have reduced pumping rates and depend strongly on microbial processing of dissolved organic matter and on the digestion of their symbionts (*27–30*). Sponge-associated microbial communities are highly host species-specific and distinct from those of the surrounding seawater (*31*, *32*). However, several studies have shown that microbial taxa associated with LMA sponges are phylogenetically more closely related to free-living microbes than those associated with HMA sponges (*33–35*), suggesting that LMA microbiomes are evolutionarily less distinct from the environmental microbial pool and may therefore be more susceptible to environmental change.

Here, we use marine sponges as a model to investigate whether host-associated microbiomes follow expectations of distributional and compositional patterns derived from those observed in free-living microbial communities. To this end, we assess differences in microbiome diversity, composition, specialization, and coevolution across hosts, while disentangling the relative importance of environmental context (shallow vs. deep) and host type (HMA vs. LMA sponges) to these differences. Our study provides a global assessment of how host-associated microbiomes are structured and have evolved across a wide range of environmental conditions and host types.

## Results

### Repeated evolution of microbiomes across host type and depth

We conducted a comparative analysis of host genetics and microbiome data from a total of 537 sponge samples, spanning ten major biogeographical areas that spread from Caribbean coral reefs ecosystems to the Antarctic and Arctic circles (Fig. 1A). This comprehensive dataset covered 200 sponge species from 23 different orders, and included 99 HMA (31 from deep and 68 from shallow waters) and 101 LMA (22 deep and 79 shallow) species (Fig. 1B, Table S1).

**Fig. 1.**
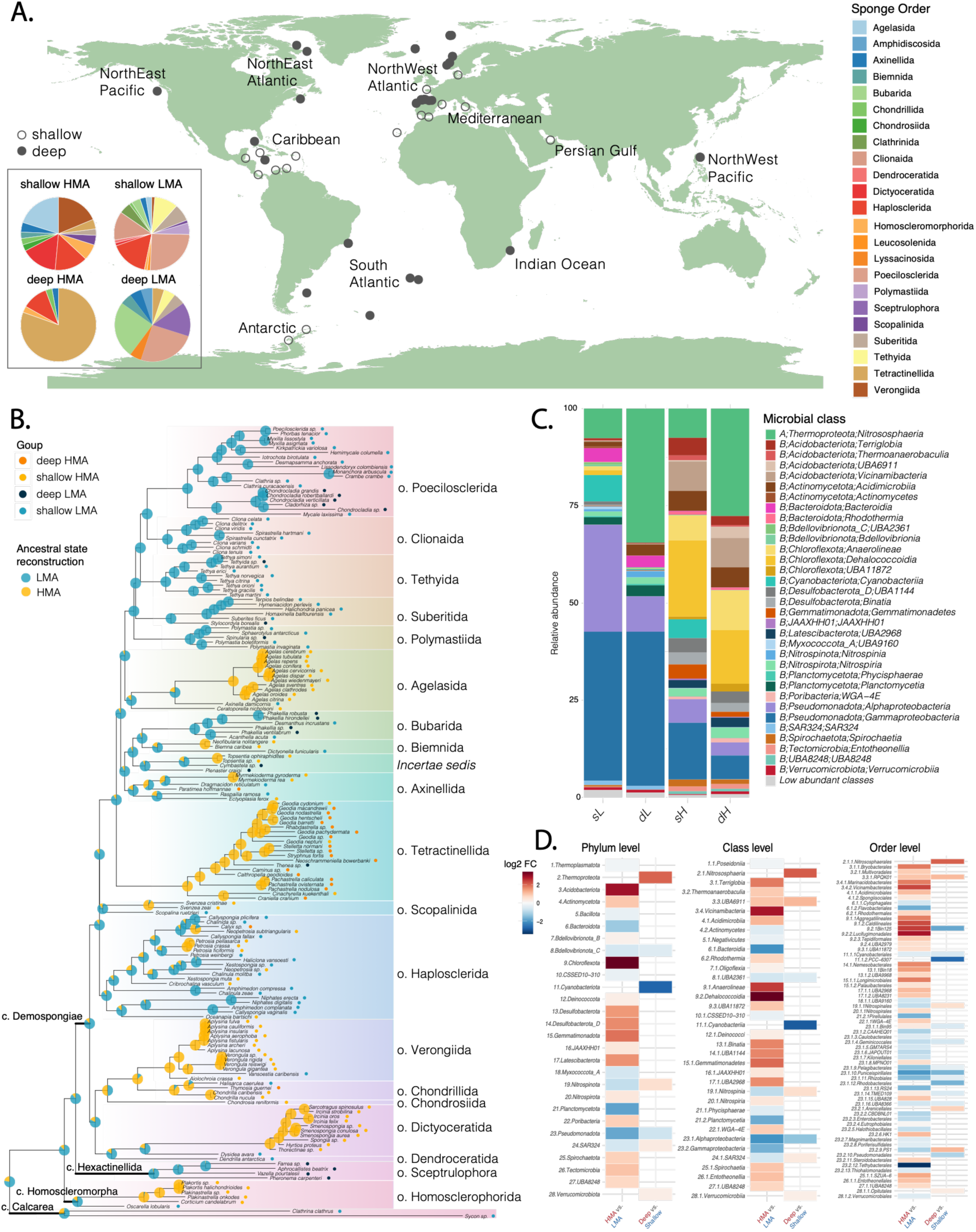
Overview of the global sponge dataset, host traits, and microbiome composition. (**A)** Map showing sampling locations, distinguishing, shallow and deep collections (circle symbols) and indicating the location of the ten major biogeographical regions covered in this study. The inset image shows the frequency distribution of sponge orders for the four sponge groups defined in this study. **(B)** Ancestral state reconstruction of HMA/LMA type across the sponge phylogenetic tree, based on Cytochrome c oxidase subunit I (COI). Sponge type and depth categories are indicated at the tips, and sponge orders are color-coded and labelled. Pie charts at internal nodes show the estimated ancestral probability of HMA and LMA type, reconstructed under an equal-rates (ER) model. **(C)** Mean relative abundance distributions of sponge-associated microbiomes at class level for each of the four sponge groups defined in this study. Classes with less than 10% relative abundance were grouped into an “Low abundant classes” category. **(D)** Differentially abundant microbial taxa identified at the phylum (left), class (middle) and order (right) levels. Color represents the log_2_ fold-change in mean abundance (HMA relative to LMA, and deep relative to shallow). Only taxa with an adjusted *P* < 0.01 (Wilcoxon test) are colored.

To examine the evolution of host type, we performed an ancestral character reconstruction on the host phylogeny based on the cytochrome oxidase I (*COI*) gene (Fig. 1B). Our results show that the HMA type evolved multiple times independently across various phylogenetic clades from LMA ancestors, as previously noted (*36*). Similarly, deep-sea sponges appeared multiple times along the phylogeny (Fig. S1), indicating independent transitions to deep environments from shallow-water lineages (*37*, *38*).

Independent evolutionary transitions repeatedly gave rise to similar microbiomes in HMA species, characterized by the dominance of phyla such as *Chloroflexota, Acidobacteriota* and *Gemmatimonadota*; whereas LMA species maintained microbiomes dominated by *Pseudomonadota, Planctomycetota* and *Bacteroidota* (Fig. 1C-D), in line with previous studies (*25*, *29*, *39–41*). In contrast to sponge type, transitions to deep environments produced only minor shifts in microbiome composition, mainly affecting relative abundances (Fig. 1C-D). Specifically, deep-sea sponge microbiomes were consistently enriched in *Thermoproteota* and the ammonia-oxidising *Nitrososphaerales* within them, whereas shallow-water microbiomes were enriched in *Cyanobacteriota* in both sponge types, and in *Alphaproteobacteria* in LMA sponges (Fig. 1D). These patterns mirror those reported for free-living microbial communities across depths (*42–44*). Ancestral state reconstruction suggested that sponge ancestral microbiomes were primarily composed of *Pseudomonadota* (*Alpha-* and *Betaproteobacteria*), and *Thermoproteota* (Fig. S2). Enrichments at lower taxonomic levels within a phylum were often not uniform (Fig. 1D) and depended on the phylogenetic organization of HMA- and LMA-associated microbiomes (see Supplementary Text and Fig. S3).

To further assess the evolutionary stability of sponge microbial type and depth preference following their emergence, we quantified phylogenetic signal for both binary traits using the D statistic (*45*). Both traits exhibited significant phylogenetic signal (p < 0.001), with D values below zero, indicating stronger phylogenetic clustering than expected under a Brownian threshold model. Sponge type (*D* = -0.233) showed a stronger phylogenetic signal than depth (*D* = -0.065), indicating that sponge type is more evolutionarily conserved than depth preference.

### More diverse yet more similar symbionts in deep-HMA species

Depth affected HMA and LMA microbiomes differently, as shown by a significant host type x depth interaction (R² = 0.03, F₁,₁₉₆ = 8.5, p ≤ 0.001), in addition to the main effects of host type (R² = 0.13, F₁,₁₉₆ = 31.9) and depth (R² = 0.05, F₁,₁₉₆ = 11.9; both p ≤ 0.001; Table S2). Differences between groups, however, may partially result from different dispersions within-group (betadisper, p ≤ 0.001 for all factors). Pairwise comparisons showed that depth explained considerably more variance in HMA (R² = 0.17, F = 19.6) than in LMA sponges (R² = 0.04, F = 4.4), and that differences between HMA and LMA communities were more pronounced in deep (R² = 0.27) than in shallow habitats (R² = 0.14; all p ≤ 0.001, BH-adjusted, Table S2). These patterns were evident in the PCoA, which showed a clear separation between HMA and LMA microbiomes along the first axis, with depth structuring HMA samples along the second axis, whereas LMA samples were only weakly separated by depth (Fig. 2A). Beyond host type and depth, sponge order explained the largest proportion of variance (R² = 0.32, F₂₂,₁₇₇ = 3.82, p ≤ 0.001), consistent with previous studies (*32*, *46*, *47*), and geographic location explained 13% (R² = 0.13, F₉,₁₉₀ = 3.07, p ≤ 0.001; Table S2). However, neither order nor location produced evident clustering in the PCoA or the dendrogram (Fig. 2B-C), with the exception of the order Agelasida, which formed a distinct branch separate from the main HMA and LMA clades in the dendrogram (see Supplementary Text for details on the Agelasida case).

**Fig. 2.**
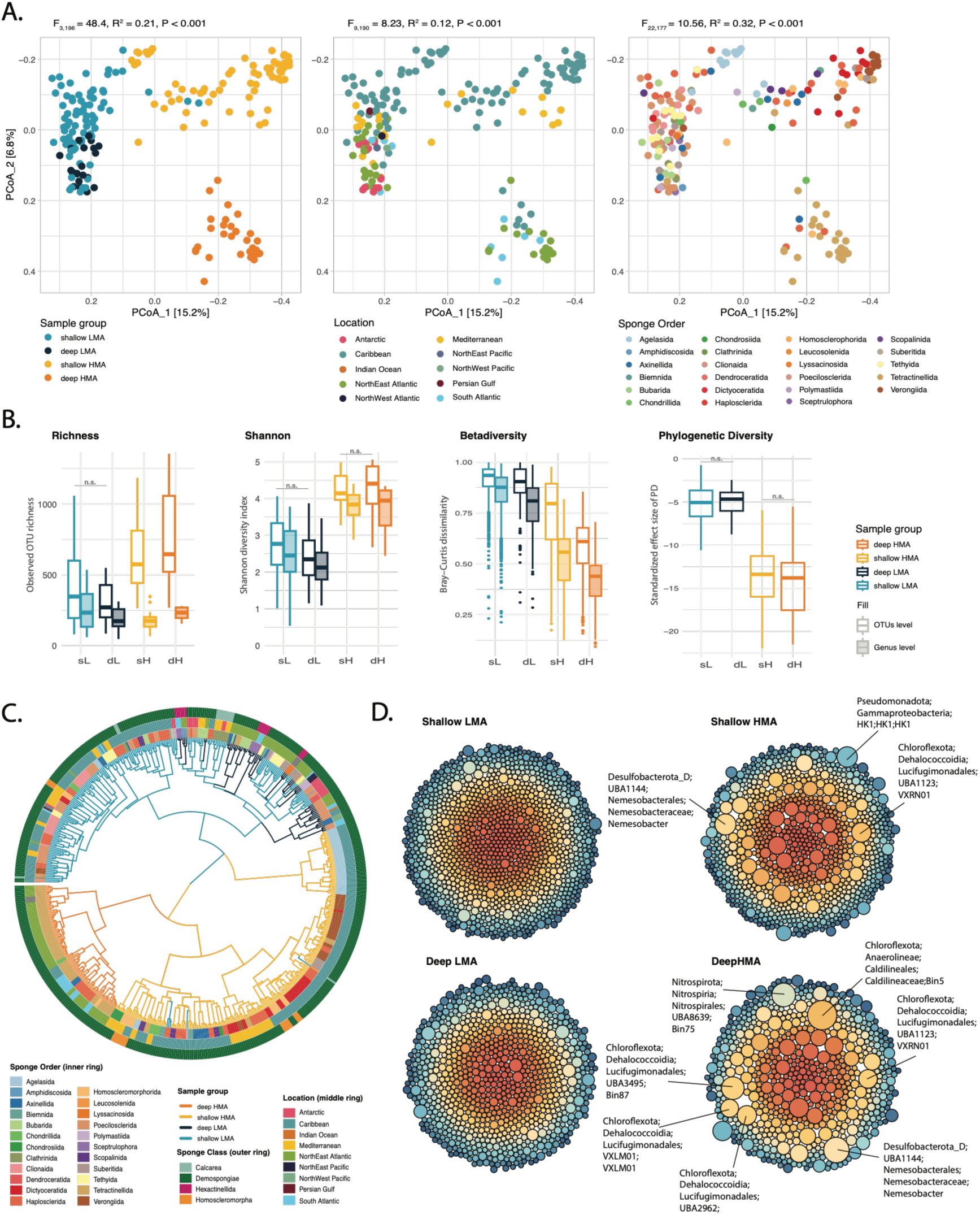
Diversity and taxonomic structure of sponge microbiome communities across host type and depth. **(A)** Principal Coordinate Analysis (PCoA) plots based on Bray-Curtis dissimilarities of sponge microbiome OTU communities, with samples colored according to group, sampling location, and sponge order. **(B)** Alpha- and Beta-diversity metrics of sponge microbiomes across groups including richness, Shannon diversity index, Bray-Curtis community dissimilarity, and standardized effect size of phylogenetic diversity (SES.PD). **(C)** Circular dendrogram based on Bray-Curtis dissimilarities of sponge microbiome OTU communities. Branches are colored by group, and rings indicate sponge order (inner), sampling location (middle), and sponge class (outer). **(D)** Microbiome composition at genus level visualized using circle packing plots. Each circle represents the mean frequency of microbial OTUs within a genus. Some of the microbial genera showing the largest expansion in HMA species are labelled by their taxonomy.

Diversity measures at the OTU level showed that HMA communities were richer, more diverse, more even, and more similar to each other than LMA communities at both depths (generalized least squares models with group-specific variances, all p < 0.001; Fig. 2B, Table S3). This pattern reflected the strong dominance structure of LMA microbiomes, which were often dominated by one or two OTUs accounting for about half of the relative abundance (Fig. S4), resulting in highly uneven communities (Fig. S5). Depth, however, affected richness and community homogeneity differently in each sponge type (host type × depth interactions, p = 0.011 and p = 0.003, respectively; Table S3). Deep-HMA sponges contained a richer symbiotic

community than their shallow-water counterparts (+21%, p = 0.039), yet symbionts were more similar across species in deep sponges (i.e., median pairwise Bray-Curtis dissimilarity of 0.61 *vs*. 0.80 in deep *vs* shallow; distance to centroid, p < 0.001; Fig. 2B). Shannon diversity, however, did not differ between deep and shallow-water HMA sponges (p = 0.859; Fig. 2B). In contrast, microbiomes from LMA sponges showed similar richness and diversity between shallow and deep environments (p=0.095 and p=0.077, respectively), although deep-LMA sponges hosted slightly more similar communities than their shallow counterparts (median of 0.90 *vs*. 0.93 in deep *vs* shallow; distance to centroid, p = 0.046). Together, these results contrast with the previous notion that deep-sea sponge microbiome communities are less complex (i.e. lower richness) than those of shallow-water species (*46*), at least for HMA sponges. This discrepancy may stem from the exclusion of the archaeal community in previous studies, as archaea are dominant in many deep-sea species. Despite their higher taxonomic diversity, HMA communities had lower phylogenetic diversity than expected under the null model at both depths, indicating that they contain closely related microbes (Fig. 2B).

Consistent with the strong community homogeneity observed among deep-HMA sponges, an occupancy analysis showed that they harbored a higher proportion of core symbionts (i.e., OTUs present in >70% of host species) and fewer specialists than deep-LMA sponges and shallow sponges of either type (Fig. S6; Supplementary Text). Regarding OTU overlap between groups, 14.3% of OTUs were shared between LMA and HMA species (representing 24.8 and 25.3% of the OTUs in each group, respectively), and this overlap was even smaller in deep-sea habitats. In contrast, 52% of OTUs from deep sponges were shared with shallow sponges, and this proportion reached 57.2 % in deep-LMA sponges (Fig. S7; Supplementary Text).

Compared to the analysis at the OTU level, richness and Shannon diversity at the genus level declined markedly in HMA species, whereas the decline was less pronounced in LMA species (Fig. 2B). HMA species are thus characterized by extensive diversification within several microbial genera, especially those belonging to the phylum *Chloroflexota*, while LMA microbiomes host more genera with fewer OTUs each (Fig. 2D). This suggests that HMA species harbor a high diversity of closely related microbial taxa, potentially reflecting recent diversification within the host, in line with their lower phylogenetic diversity. Notably, this expansion of specific genera in HMA sponges did not correlate with higher relative abundances of those genera.

### Limited coevolution despite strong phylosymbiosis in deep environments

Phylosymbiosis reflects the extent to which phylogenetically similar host species harbor more similar symbiont communities than distantly related ones (*48–50*). Previous studies have shown phylosymbiosis occurring only in HMA sponges, but this has been tested only on shallow water species (*32*, *36*, *51*, *52*). Our global dataset analysis confirmed this pattern and further revealed that the phylosymbiotic signal (corrected by geographic distance) remarkably strengthens with depth in HMA sponges (partial Mantel’s *r* = 0.257 in shallow, and *r* = 0.433 in deep environments; Fig. 3A). Unexpectedly, while shallow-water LMA species did not exhibit evidence of phylosymbiosis (partial Mantel’s *r* = -0.093), a strong signal emerged in deep-water LMA sponges (partial Mantel’s *r* = 0.330). Microbial dissimilarity also increased with geographic distance in all groups except shallow-HMA (Fig. S8), but geographic and phylogenetic distances were themselves not correlated (*r* ≈ 0, *p* > 0.05), ruling out their collinearity. This indicates that microbiome assembly is governed by host phylogeny, but also independently affected by a geographical component.

**Fig. 3.**
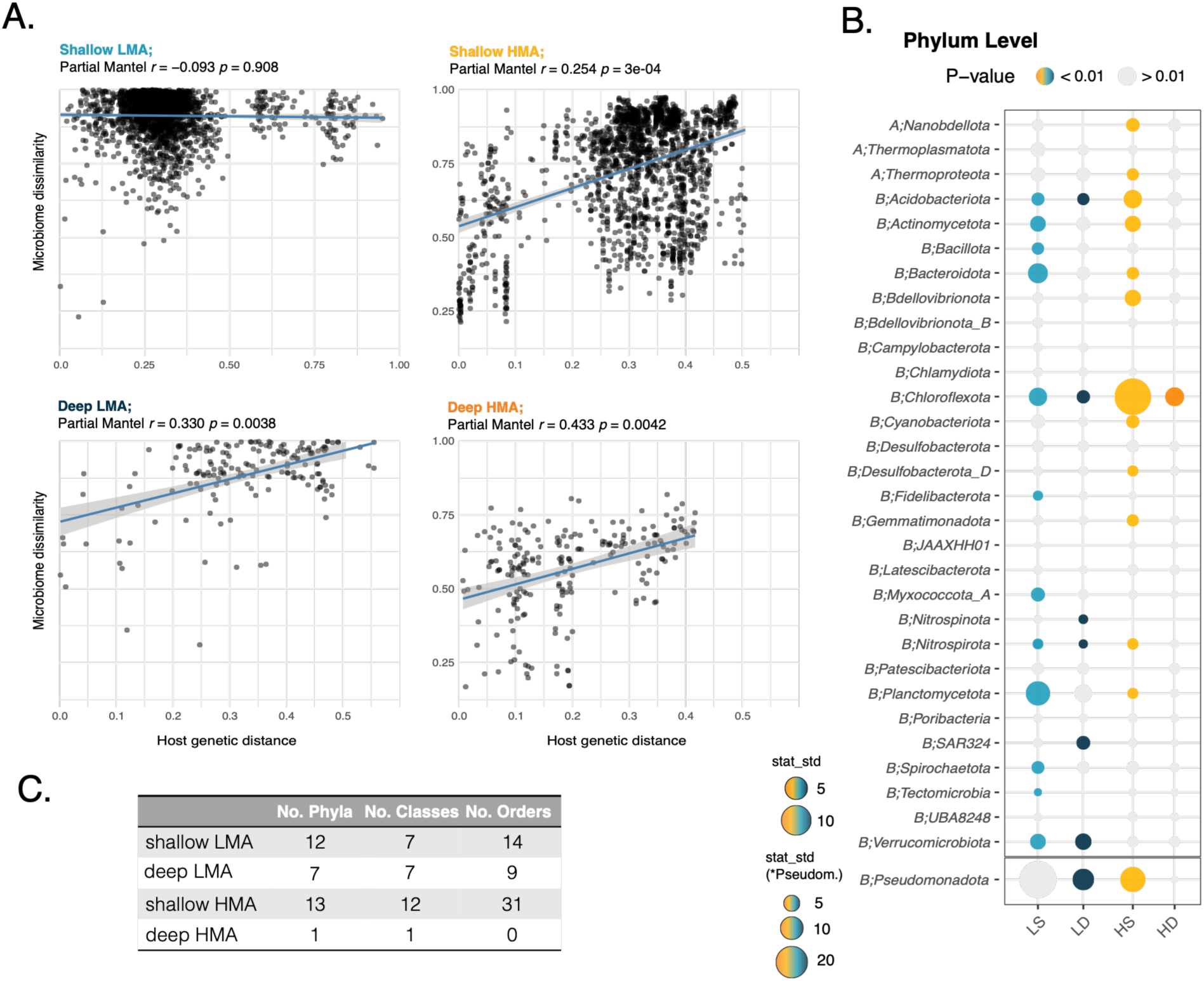
Host-microbiome evolutionary dynamics. **(A)** Correlation between microbiome dissimilarity (measure using Bray-Curtis dissimilarity of OTU composition across samples) and host genetic distance (pairwise phylogenetic distances from the constructed sponge tree) across the four sample groups. Partial Mantel test statistics (Pearson’s r) and p-values (p) are indicated above each plot; lines show linear regressions with standard error (SE). (**B)** Cophylogeny analysis between hosts and symbionts, assessed at the microbial phylum level. Bubble size represents the standardized ParaFit test statistic. Colored bubbles indicate statistically significant associations (p < 0.01), and non-significant associations are shown in grey. Phylum *Pseudomonadota* is separated by a line, as its ParaFit values are shown separately below. A = Archaea; B = Bacteria. **(C)** Summary of significant microbial lineages at phylum, class, and order levels for each group; visualizations are provided in the Fig. S10.

Phylosymbiosis most often arises from the influence of ecological factors, including the environmental acquisition of microbes contingent on host physiology, habitat and geographical preferences (*26*, *48*, *53–55*), but could also be due to coevolution or codiversification, a process characterized by reciprocal adaptations and selection between hosts and microbial symbionts. To evaluate whether, and to what extent, the observed phylosymbiosis emerges from codiversification, we employed the ParaFit statistic (*56*), a widely used method for assessing congruence between host and symbiont phylogenies (i.e., co-phylogeny analyses). The microbial taxonomic level at which co-phylogenetic signal can be detected is still poorly understood, with previous studies looking at the family level in corals (*57*) or below class-level in sponges, specifically for the class *Dehalococcoidia* (*36*). Since we detected enrichments at different taxonomic levels (Fig. 1D), co-phylogeny analyses were performed separately for each prokaryotic phylum, class and order (Fig. 3B-C, and Fig. S9). If coevolution were a major driver of phylosymbiosis, we would expect stronger co-phylogenetic signals in HMA than in LMA hosts and in deep compared with shallow environments, consistent with the higher phylosymbiotic signal observed in HMA species and deep habitats. However, this was not the case. Deep-water HMA species showed co-phylogeny only with phylum *Chloroflexota*, whereas deep-LMA species showed significant co-phylogeny with seven microbial phyla and nine orders (Fig. 3B, Fig. S9). Co-phylogeny was, however, much more pronounced in shallow sponges, with HMA species showing significant signals with 13 phyla and 31 orders, particularly *Chloroflexota* and several of its constituent orders. Similarly, shallow-LMA species exhibited strong co-phylogenetic signals for 12 phyla and 14 orders (Fig. 3B-C). Notably, this strong co-phylogenetic signal in shallow-LMA sponges contrasts with their comparatively weak phylosymbiosis, suggesting that the evolutionary congruence of individual host-microbe associations does not necessarily generate a community-wide phylosymbiotic pattern.

## Discussion

Our results show that host-associated microbial diversity does not always respond to depth in the same way as planktonic microbial communities in the ocean, with host type emerging as the primary driver of this divergence. In planktonic microbial communities, richness increases with depth (*58–62*), while at the same time communities become more compositionally homogeneous (*6*, *9*). In our study, only HMA sponge microbiomes mirrored this trend. LMA sponges, in contrast, consistently harbored less diverse microbiomes that remained highly distinct among host species across depths. Moreover, deep and shallow LMA microbiomes were more similar to each other than were deep and shallow HMA microbiomes. Together, these results suggest that the environmental stability and connectivity of deep waters may shape microbiome diversity more strongly in HMA than in LMA sponges, despite LMA microbiomes being widely regarded as more similar to those of the surrounding seawater (*33–35*).

Results from ancestral state reconstruction together with analyses of taxonomic composition and phylogenetic diversity, suggest that HMA sponges independently acquired microbial lineages from the same phyla, which subsequently diversified within the host, potentially favoured by the HMA body plan, which is suited to hosting tightly packed microbial communities (*28*). This diversification appears to have occurred preferentially in specific lineages (e.g., *Chloroflexota*; Fig. 2D), resulting in communities of closely related taxa (i.e., high taxonomic diversity but low phylogenetic diversity; Fig. 2B). This pattern of diversification may reflect the relatively stable internal environment of the sponge and reduced competition for resources compared with free-living communities. Such in-host diversification could explain why symbionts of HMA sponges show greater phylogenetic distinctiveness from their free-living relatives than do the lineages associated with LMA sponges, as reported in other studies (*34*, *35*, *63*, *64*). By contrast, LMA-associated microbes may be acquired repeatedly from the surrounding seawater, with limited opportunity for in-host diversification, resulting in communities that, although compositionally distinct from those in the water column, retain greater phylogenetic similarity to their environmental source lineages.

Phylosymbiosis signal was strongest in deep environments for both HMA and LMA species. In shallow-water HMA sponges the signal remained strong, whereas in shallow-water LMA sponges microbial compositional dissimilarity was consistently high and did not correlate with host genetic distance. Planktonic microbial communities in shallow, coastal waters are known to be highly heterogeneous and spatially patchy, reflecting strong environmental variability and limited connectivity (*9*). Because LMA sponges show a reduced dependence on symbiont-derived nutrition and pump larger volumes of seawater (*29*, *30*, *65*, *66*), high environmental heterogeneity likely contributes to the reduced phylogenetic structuring of their microbiomes. We hypothesise that microbiomes of LMA sponges may emerge from a more flexible assembly process, whose outcome depends strongly on the availability of suitable symbionts in the environment and on the host’s need for them. In deep environments, where the microbial pool is more homogeneous (*6*, *9*), and conditions are more stable, LMA sponges could select their preferred symbionts, thereby reinforcing phylosymbiosis. By contrast, greater environmental heterogeneity and patchier microbial distributions in shallow waters (*59*) may limit the availability of suitable symbionts, leading LMA sponges to recruit functionally equivalent microbes from closely related lineages and thereby weakening phylosymbiosis. Moreover, accumulating locally adapted microorganisms from the seawater could reduce the vulnerability of LMA sponges to environmental perturbations (*64*). However, HMA sponges, with lower filtration rates and higher microbial dependency (*28*), may exert stronger selection, being capable of maintaining more consistent associations with their chosen microbial partners across environments and thereby preserving a strong phylosymbiotic signal even in shallow habitats.

We tested whether phylosymbiosis was linked to cophylogenetic processes and found a striking decoupling between the two. Coevolutionary signals were largely restricted to shallow environments, and declined with depth despite phylosymbiosis itself increasing. A plausible explanation would be the dominance of homogenising eco-evolutionary processes in deep waters. For coevolution to happen several conditions need to be satisfied. The most important one is environmental heterogeneity combined with spatial isolation, allowing different species to interact and evolve together across a mosaic-type landscape (*67*). Deep environments are characterized by the opposite, with reduced isolation, high connectivity due to high dispersal rates and environmental homogeneity likely weakening host-microbe coevolution compared to shallow systems. Consequently, phylosymbiotic patterns, where closely related hosts harbor more similar symbiont assemblages, may, in shallow environments, arise from genuine shared coevolutionary history of hosts and symbionts, where closely related symbionts inhabit closely related hosts. In deep environments, by contrast, phylosymbiosis is likely driven by different underlying processes, including differential filtering of microbial communities by host traits, host phylogeography, and environmental effects on microbiome assembly (*49*, *55*, *68*), or a mix of shared evolutionary history and host filtering (*69*).

Our study reveals the ecological and evolutionary processes that underpin the adaptation of host-associated microbiomes to deep-sea environments. Environmental constraints operating in shallow and deep habitats affect host sponges differently, reflecting differences in microbiome diversity and host selective capacities. Evolutionary dynamics also differed: in stable and homogeneous environments like the deep-sea, strong phylosymbiosis did not emerge from coevolutionary dynamics, but from host filtering and environmental effects, whereas in heterogeneous shallow environments, microbiome specificity results from tighter coevolutionary dynamics.

## Materials and Methods

### Dataset compilation

A comprehensive dataset was assembled by combining sequences from newly sampled sponge species collected during multiple campaigns with publicly available datasets generated with the same primer pair and retrieved from the Sequence Read Archive (SRA) of the National Center for Biotechnology Information (NCBI). The final compilation included 61 new species, together with data from six previously published studies by the authors (*70–75*) and two additional studies (*36*, *76*) (Table S1).

Although the datasets contained numerous replicates for some species, we limited the analysis to a maximum of four replicates per sponge species and sampling location to reduce dataset size and ensure a balanced design. Sixty-two species were represented by fewer than four replicates. In total, the dataset comprised 537 samples representing 200 sponge species across 24 orders. These sponges were collected from six major geographic regions worldwide (Fig. 1A). Samples collected by scuba diving at depths shallower than 30 m were classified as *shallow*, corresponding to waters above the deep chlorophyll maximum (*9*). Samples collected between 94 and 4,196 m depth were classified as *deep*. Sponge species were assigned to either the high microbial abundance (HMA) or low microbial abundance (LMA) category following the classification reported in the original studies from which the data were obtained, based on previous reports when no classification was available (*25*), or, for newly sampled species, based on TEM and 16S rRNA gene sequence data (*41*). This resulted in four host groups: 68 shallow HMA species, 31 deep HMA species, 79 shallow LMA species, and 22 deep LMA species.

Some geographic regions, such as the Caribbean and the Northwest Atlantic, contained representatives of all four groups (Fig. 1A). Likewise, each group encompassed multiple sponge orders. The taxonomically least diverse group was the deep-HMA, which comprised five orders and was dominated by *Tetractinellida* (29 species). This taxonomic composition reflects the characteristic sponge fauna of the North Atlantic and boreal deep-sea habitats, as previously described (*77*, *78*). In fact, *Tetractinellida* is the second most diverse sponge order globally and is widespread and particularly successful in deep-sea environments (*79*, *80*). Nevertheless, this taxonomic bias is unlikely to influence our analyses, as the deep-HMA group still spans a substantial range of host genetic distances (maximum pairwise distance = 0.42), comparable to that of shallow HMA species (0.50), and includes samples collected from all major deep-sea regions represented in this study.

### Sequencing processing and taxonomic assignment

Community composition was studied by sequencing the V4 hypervariable region of the 16S rRNA gene of prokaryotes. For newly processed samples, DNA was extracted from about 20 mg of tissue preserved in ethanol using the DNeasy Blood and Tissue kit (Qiagen). The V4 region was sequenced using primers 515F-Y (5’ GTGYCAGCMGCCGCGGTAA 3’) (*81*) and 806RB (5’ GGACTACNVGGGTWTCTAAT 3’) (*82*), targeting both archaea and bacteria. DNA preparation, read processing, and taxonomic assignment were performed following previous protocols (*74*). Briefly, DNA amplification was done in duplicates with 28 PCR cycles, libraries prepared with the Nextera XT DNA Library Preparation Kit (Illumina Inc.), and next-generation, paired-end sequencing was performed at the Complutense University of Madrid on a MiSeq System (Illumina) using v3 chemistry (2 × 300 bp).

Our newly generated sequences and those retrieved from public repositories were combined into a single dataset and processed jointly to allow direct comparisons. Read processing followed the MiSeq SOP protocol (*83*) implemented in Mothur v1.48.0 (*84*). Because the dataset integrates samples generated across multiple studies, laboratories, and sequencing runs, sequences were clustered into operational taxonomic units (OTUs) at 97% sequence similarity rather than resolved as amplicon sequence variants (ASVs), to minimize the impact of technical variation among datasets that could otherwise be interpreted as biological differences. Furthermore, recent analyses based on high-quality genomes from the Genome Taxonomy Database (GTDB) (*85*) have shown that intragenomic variation in the microbial 16S rRNA gene can reach 0.0306 sequence dissimilarity (*86*). Chimeric sequences were identified and removed using the *chimera.uchime* command in Mothur. The dataset was then rarefied to the smallest library size (9,034 reads per sample), and then filtered to remove any OTU with less than 10 counts across all samples. The resulting dataset comprised 9,880 OTUs. Finally, community composition was averaged across replicates from the same sponge species, yielding a single mean relative abundance value for each OTU per species.

Classification of OTUs was performed using the GTDB taxonomy release 232 (15^th^ of April 2026). This release comprises 901,341 genomes; however, not all genomes contain annotated 16S rRNA genes, and many genomes contain multiple copies of this gene. To generate a curated reference database, we retained genomes containing 16S rRNA sequences longer than 1,000 bp, with a single SSU sequence, and excluded genomes annotated at the lowest-quality assembly level (i.e., *contig*) (*86*). We also used the publicly available GTDB bacterial and archaeal reference trees (bac120_r232.tree with 189,801 tips, and ar53_r232.tree with 10,122 tips) as our microbial phylogenetic backbone, rather than constructing independent local phylogenies based solely on our amplicon sequences, as these short fragments (∼290 bp) provide limited phylogenetic resolution (*87*). The GTDB trees were inferred from concatenated alignments of 120 and 53 conserved proteins for Bacteria and Archaea, respectively, and represent the evolutionary relationships among GTDB representative genomes. We subsequently pruned these trees to retain only the curated genomes containing the selected 16S rRNA sequences, resulting in a reference tree of 39,159 tips. This backbone tree was used for phylogenetic placement of OTUs using the Evolutionary Placement Algorithm (EPA-ng) (*88*). Phylogenetic placement provides a more accurate framework for taxon identification than sequence similarity approaches (*88*, *89*). Downstream processing and taxonomic assignment were performed using *gappa* v0.9.0 (*90*).

### Host phylogenetic reconstruction

Cytochrome c oxidase subunit I (COI) sequences of 164 species in our dataset were downloaded from NCBI (Table S1). In addition, twelve COI sequences were generated from our stored specimens using the Folmer primers as in Riesgo et al. (*91*). The final dataset included 167 COI sequences from the 200 species analysed in this study. Consequently, all analyses based on the host phylogeny were restricted to the 167 species with available COI sequences.

COI sequences were then aligned separately using MAFFT (*92*) with the default parameters. The COI alignment was trimmed to 603 nucleotides. Phylogenetic reconstruction was performed using Maximum Likelihood (ML) method in RAxML-NG v. 1.1.0 (*93*) implemented in raxmlGUI v.2.0 (*94*), after selecting for the most likely nucleotide substitution model using jModelTest (*95*), and setting the search strategy to ML + thorough bootstrap + consensus. The final evolutionary model was GTR+FO, with 100 bootstraps replicates and 10 starting trees. Because the COI cannot resolve order-level relationships, we used the order-level topology of the current accepted phylogenetic backbone of sponges (*96*, *97*) to constrain the phylogenetic reconstruction. Pairwise phylogenetic distances from the constructed sponge tree were calculated using *cophenetic.phylo* from the R package *ape v. 5.8.1* (*98*)

Ancestral character state reconstruction of HMA and LMA status was performed using maximum likelihood with the ace function in the R package ape. Discrete traits were analysed under an equal-rates (ER) Markov model, which assumes a single transition rate between character states. The likelihoods of ancestral HMA and LMA states were estimated at all internal nodes of the phylogeny.

To reconstruct ancestral microbiome composition, microbial abundances were aggregated at microbial class level for each sponge species. Abundance data were converted to proportions, (*98*) and zero values were replaced with a small pseudocount (10⁻⁶). Because microbiome data are compositional, proportional abundances were transformed using an isometric log-ratio (ILR) transformation implemented in the R package compositions. Each ILR coordinate was independently reconstructed at internal nodes using maximum-likelihood ancestral-state reconstruction under a Brownian-motion (BM) model, implemented in the *anc.ML* function of *phytools* (*99*). The reconstructed ancestral ILR coordinates were subsequently back-transformed using the inverse ILR transformation to obtain estimated ancestral microbial proportions. For visualization, ancestral and extant microbiome compositions were represented by the 33 most abundant microbial taxa across the dataset, with all remaining taxa grouped as “Other”.

Phylogenetic signal (PS) for sponge status (HMA/LMA) and habitat (shallow/deep) was assessed using the *D* statistic (*45*), as implemented in the *phylo.d* function of the R package *caper*. The D statistic quantifies the strength of phylogenetic signal in binary traits. A value of D = 1 indicates that the observed trait is randomly distributed across the phylogeny, whereas D = 0 indicates a level of phylogenetic clustering consistent with evolution under a Brownian motion model. Values of D < 1 indicate increasing phylogenetic clustering, while D > 1 suggests phylogenetic overdispersion (*45*).

### Microbial composition analysis

Alpha diversity metrics, including richness and Shannon diversity, were calculated from rarefied OTU tables for each sample and averaged per host species. Differences among groups were assessed using generalized least squares (GLS) models with host type, depth and their interaction as fixed effects, allowing for group-specific variances (varIdent) to account for heterogeneity of variances among groups (*nlme* package). Richness was log-transformed prior to analysis. Pairwise contrasts between depths within each host type, and between host types within each depth, were computed with the *emmeans* package using Satterthwaite degrees of freedom. Phylogenetic diversity (PD) was estimated as the standardized effect size (SES) of Faith’s PD, which quantifies phylogenetic diversity relative to a null expectation (*100*, *101*). This metric was based on OTU presence–absence and the microbial phylogenetic tree, and was calculated with the ses.pd function of the *picante* package. SES.PD values were analyzed with the same GLS framework.

For beta diversity analysis, relative abundances from rarefied OTU tables were log2 transformed prior to calculating Bray–Curtis dissimilarities. Beta diversity within groups was visualized as pairwise Bray–Curtis dissimilarities among host species. Because pairwise values are not independent, differences among groups were tested on distances to group centroids (betadisper, vegan) (*102*), calculated for the four host type x depth groups and compared using the same GLS framework as for alpha diversity.

Differences in community structure among groups were visualized using principal coordinates analysis (PCoA) based on Bray–Curtis distances, implemented with the *cmdscale* function in the R package stats. Statistical differences among groups were assessed using permutational multivariate analysis of variance (PERMANOVA) with the *adonis2* function in the R package vegan (v. 2.6-4), including host type, depth and their interaction, followed by pairwise PERMANOVAs between depths within each host type and between host types within each depth, with p-values adjusted using the Benjamini–Hochberg method. Homogeneity of multivariate dispersions, an assumption of PERMANOVA, was tested for each factor with betadisper and permutation tests (permutest, 999 permutations).

Differences in microbial relative abundances among host types and depth categories were assessed using Wilcoxon rank-sum tests at three taxonomic levels: phylum, class, and order (with p-values adjusted using the Benjamini–Hochberg method). The distribution of OTUs among microbial genera and across sample groups was visualized using circle-packing plots, where each circle represents the average number of microbial OTUs assigned to a genus.

To characterize core, shared and specialist microbial features, OTUs representing less than 0.001 % relative abundance per sample were removed by setting their abundance to zero, thereby reducing the influence of extremely rare taxa. Within each sample group, OTUs were classified as specialist if they were detected in only one sponge species, as shared if they occurred in more than one sponge species, and as core if they were present in >70% of sponge species within the group. This threshold is an arbitrary, but was chosen given the large number of species considered (n = 200); stricter thresholds (≥90%) yielded few core taxa, and changing core definitions has been shown not to substantially affect the results for closely related samples (*103*). The core definition was not used for further diversity or statistical analysis. These categories were summarized both as the frequency of occurrence across host species and as their contribution to total relative abundance. Between sample groups, generalist OTUs were defined as those occurring in both groups, whereas endemic OTUs were restricted to a single group.

#### Phylosymbiosis and cophylogeny

Phylosymbiosis between microbial community composition (Bray-Curtis dissimilarity) and host genetic distance (cophenetic distances) was assessed using partial Mantel tests, controlling for geographic distance. These geographic distances were calculated as oceanic least-coast distances (i.e., shortest water route between two marine locations) using the *rnaturalearth* R package and Natural Earth data at a spatial resolution of 0.1° (∼11 km), and the *gdistance* R package to calculate pairwise distances in km. The partial Mantel approach evaluates the correlation between two distance matrices using Pearson correlation coefficients, controlling for the effect of a third matrix, with statistical significance determined through 999 permutations of the distance matrices.

Cophylogenetic associations between hosts and microbial communities were assessed using the permutation-based ParaFit test (*56*). This method evaluates the significance of the overall association between host and microbial phylogenies given a matrix of host–microbe presence/absence associations. The null hypothesis assumes independent evolution of hosts and microbes (*56*). Host and microbial phylogenetic trees were converted into patristic distance matrices using the *cophenetic.phylo* function from the R package *ape*. To reduce spurious associations driven by extremely rare taxa, analyses were performed using the dataset that excluded OTUs with relative abundance below 0.001%. Distance matrices were then transformed using principal coordinates analysis (PCoA) with Lingoes correction to account for negative eigenvalues. The global ParaFit test was performed using the *pf_parafit* function from the R package *ape*, based on 999 permutations, with a significance threshold of 0.01. The ParaFit test statistic was standardized by the number of associations to allow comparisons among microbial taxa.

Cophylogenetic signal was evaluated across multiple microbial taxonomic levels, including phylum, class, and order. Only datasets containing at least 10 OTUs per phylogenetic tree were retained for analysis. For each analysis, phylogenetic trees were pruned to include only the taxa present in the corresponding dataset using the *keep.tip* function from the R package *ape.* Host–microbiome associations were visualized using tanglegrams generated with the *cophylo* function from the R package *phytools* (*99*).

## Data, code, and materials availability

Raw sequence data for all sponge species studied here are available from the NCBI SRA under project PRJNA1545767. Data analysis scripts are available in the GitHub repository https://github.com/cdiezvives/phylogeny-cophylogeny-sponges.

## Acknowledgments

CDV received the support from a fellowship from “la Caixa” Foundation (ID 100010434)”, code LCF/BQ/PI22/11910040, and a Ramon y Cajal fellowship RYC2024-048180-I. MT was supported by MSCA Postdoctoral Fellowship (HORIZON-MSCA.2022-PF-01, Project 101105716). JMM was co-funded by the European Union (GA101059915 - BIOcean5D) and by the French Agence Nationale de la Recherche through LabEx TULIP (ANR-10-LABX-41).

## Notes

### Competing Interest Statement

The authors have declared no competing interest.

### Summary of Updates

The author order has been corrected.

