## Supplemental Text and Figures for "Ocean depth modifies host-microbiome ecology and evolution"

**The PDF file includes:**

Supplementary Text
Figs. S1 to S10
References

**Supplementary Text**

Phylogenetic organisation of HMA- and LMA-associated microbiomes

Enrichment analysis differed between the broad phylum level and its lower-rank taxonomic sublevels (class, order, family, etc.). For example, although the phylum Bacteroidota was generally enriched in LMA sponges, its constituent classes differed: *Bacteroidia* remained enriched in LMA sponges, whereas *Rhodothermia* was enriched in HMA sponges (Fig. 1D). These differences were underpinned by the prevalence, abundance, and phylogenetic organisation of HMA- and LMA-associated microbes within each phylum.

To analyse the phylum composition and phylogeny, we first classified microbial phyla as exclusive to one sponge status when OTUs prevalence exceeded 90%, and as mixed when prevalence was below 90% (Fig. S3). Fifteen phyla were almost entirely restricted to a single sponge status, with all or nearly all OTUs occurring exclusively in either HMA or LMA hosts (e.g. *Poribacteria*, *Planctomycetota*). Among the mixed phyla, the phylogenetic clustering of sequences was characterised using the D statistic (*1*). *Bacillota* and *Bdellovibrionata* presented D values between 0 and 1, indicating a pattern intermediate between Brownian motion and random expectation; we classified these as showing "extensive intermixing of sponge-status OTUs". The remaining phyla presented D values below 0, indicating a clumped distribution pattern, however, eight of these phyla still showed low negative values ($D > -0.25$) and were also classified as "extensive intermixing of sponge status OTUs". Phyla with $D < -0.25$ displayed a visually deeper segregation between HMA- and LMA-associated clades (e.g., *Spirochaetota, Tectomicrobia*) and were classified as "partitioned into monophyletic sponge-status clusters" (Fig. S3). These contrasting phylogenetic patterns show that the evolution of host-associated microbes has not followed a common evolutionary trajectory across bacterial phyla. While some lineages show patterns consistent with diversification after becoming associated with a particular host type, others remain phylogenetically interspersed, suggesting repeated host-type shifts or limited evolutionary differentiation.

Sponge-Coral-Specific Clusters (SCSC) are more prevalent in HMA sponges

Beyond the organization observed within our sponge sequences, we further investigated whether HMA-associated OTUs (i.e. those exclusive to or more prevalent in HMA hosts) were more closely related with the previously defined Sponge-Coral-Specific clusters (*2*, *3*) than LMA-associated OTUs. This analysis was conducted manually by inspecting the placement of our OTUs within the large curated Silva v. 138.2 SSURef NR99 phylogenetic tree, and consequently, we restricted the analysis to the 100 most abundant OTUs. Our revision revealed that all HMA-associated OTU, with the exception of Otu000012, consistently fell within a large monophyletic group composed solely of sponges or coral hosts, clearly separated from environmental samples. In contrast, LMA-associated OTUs showed a more varied pattern of associations: Approximately one-third grouped within SCCs, though typically in smaller clusters containing few other sponge species; another third clustered sponge/coral and environmental sequences; and the remaining third clustered with sediment, water, or other animal-associated sequences.

A similar pattern was previously reported for phylum *Chloroflexi*, where most sequences from HMA sponges, but only a few from LMA sponges, fell into SCC clusters (*4*); moreover, the LMA-derived *Chloroflexi* sequences clustered only with other LMA-associated bacteria (*4*). Our results suggest that this pattern extends broadly across multiple bacterial phyla associated with HMA sponges, however, our LMA-associated not only clustered with other LMA-associated sequences but also with HMA-derived bacteria.

Among Archaea, the cosmopolitan Otu000001 was placed in a mixed clade with sponges and environmental samples. Other HMA-associated archaea were also not clearly placed within SSC monophyletic clusters, suggesting either a more relaxed association with sponges or an underrepresentation of sponge-derived archaea in the reference tree.

Deep and shallow hosts share a large proportion of symbionts

Core OTUs were more numerous and abundant among HMA than LMA species, reaching higher proportions with depth in both host statuses (HMA: shallow 22.3 ± 8.1% and deep 63.8 ± 13.1% relative abundance (RA); LMA: shallow 10.6 ± 13.2% and deep 19.9 ± 15.6% RA; Fig. S6). In LMA species, core OTUs were frequently dominated by a single and highly abundant archaeal OTU (Otu00001) belonging to the genus *Nitrosopumilus*. Regarding specialist OTUs (i.e., unique to a sponge species), these were more frequent and abundant in LMA than HMA species, decreasing slightly with depth (HMA: shallow 5.1 ± 8.6% and deep 2.05 ± 4.4 % RA; LMA: shallow 23.2 ± 23.6 % RA and deep 21.2 ± 28.3% RA; Fig. S6), and were distributed across all microbial phyla.

Across groups, LMA and HMA species shared approximately 14.3% of OTUs overall, which represented ~25% of OTUs within each category (Fig. S7A). This proportion decreased in deep environments, where only 9.7-14.5 % of OTUs were shared per group, suggesting greater microbiome divergence between sponge statuses with increasing depth. In contrast, comparisons across depth categories revealed a larger proportion of generalists OTUs, with 21.4% shared overall, representing up to 52% of deep OTUs and 26.7% of shallow OTUs. This proportion increased further in deep-LMA species, reaching 57.1%. Shared OTUs were generally abundant, accounting for a substantial fraction of the mean relative abundance per group (Fig. S7B).

This large overlap of OTUs between deep and shallow hosts can be explained by two non-mutually exclusive scenarios involving connectivity between environments and microbiome transmission through generations. If deep- and shallow-water microbial communities are vertically stratified (*5*, *6*), the shared OTUs must have been largely retained through vertical transmission from a common ancestral microbiome. Alternatively, if depth imposes little constraints on microbial dispersal Chen 2024 (*7*, *8*), then horizontal acquisition, which is known to play a major role in shaping host-specific microbiomes (*9*, *10*), could recruit similar microbial taxa from a well-connected microbial pool, without necessarily requiring faithful vertical transmission.

The Agelasida case

Agelas stood as a different order of all other species. In the PCoA analysis formed a differentiated group (Fig 2A). In the dendrograms formed a distinct independent branch separated from HMA and LMA species (Fig. 2C). This oddity has been pointed out before, because although *Agelas* spp. are classified as HMA sponges based on diversity and phylogenetic assignment of their microbiomes, their microbial communities are less dense than LMA species (*11*) , and clustered between HMA and LMA species (*12*, *13*).

Notably, species within the order Agelasida formed specific host-microbiome associations with distinct microbial OTUs (Fig. S10 A-B), which were phylogenetically more distant and specific of the Agelasida order in some phyla (Fig. S10 C-D). This has also been observed in other species of the genus *Agelas* from mesophotic reefs (*14*).

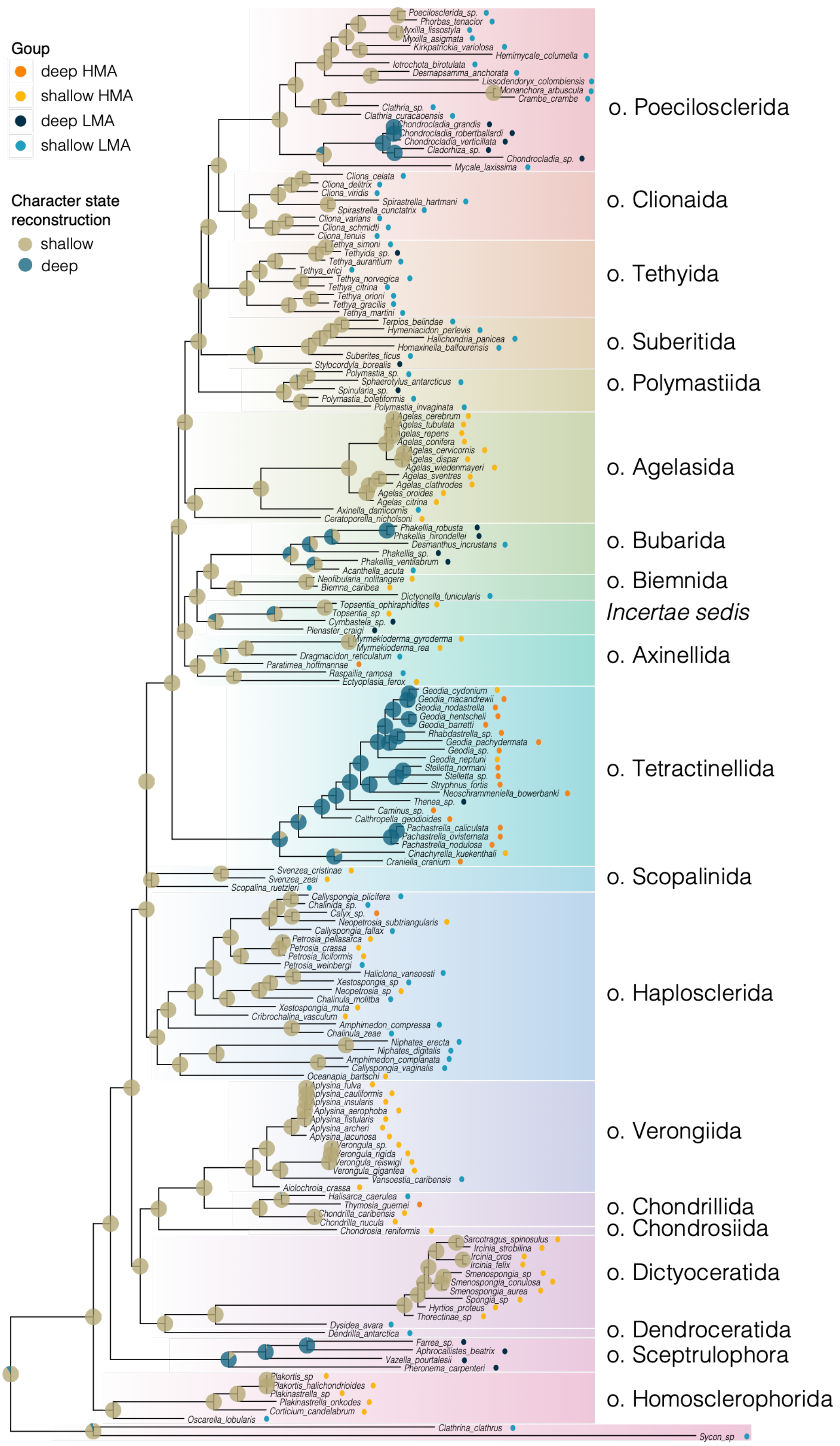


**Fig. S1. Ancestral state reconstruction of shallow and deep preference across the sponge phylogenetic tree, based on Cytochrome c oxidase subunit I (COI).** Sponge status and depth categories are indicated at the tips, and sponge orders are color-coded and labelled. Pie charts at internal nodes show the estimated ancestral probability of HMA and LMA status, reconstructed under an equal-rates (ER) model.

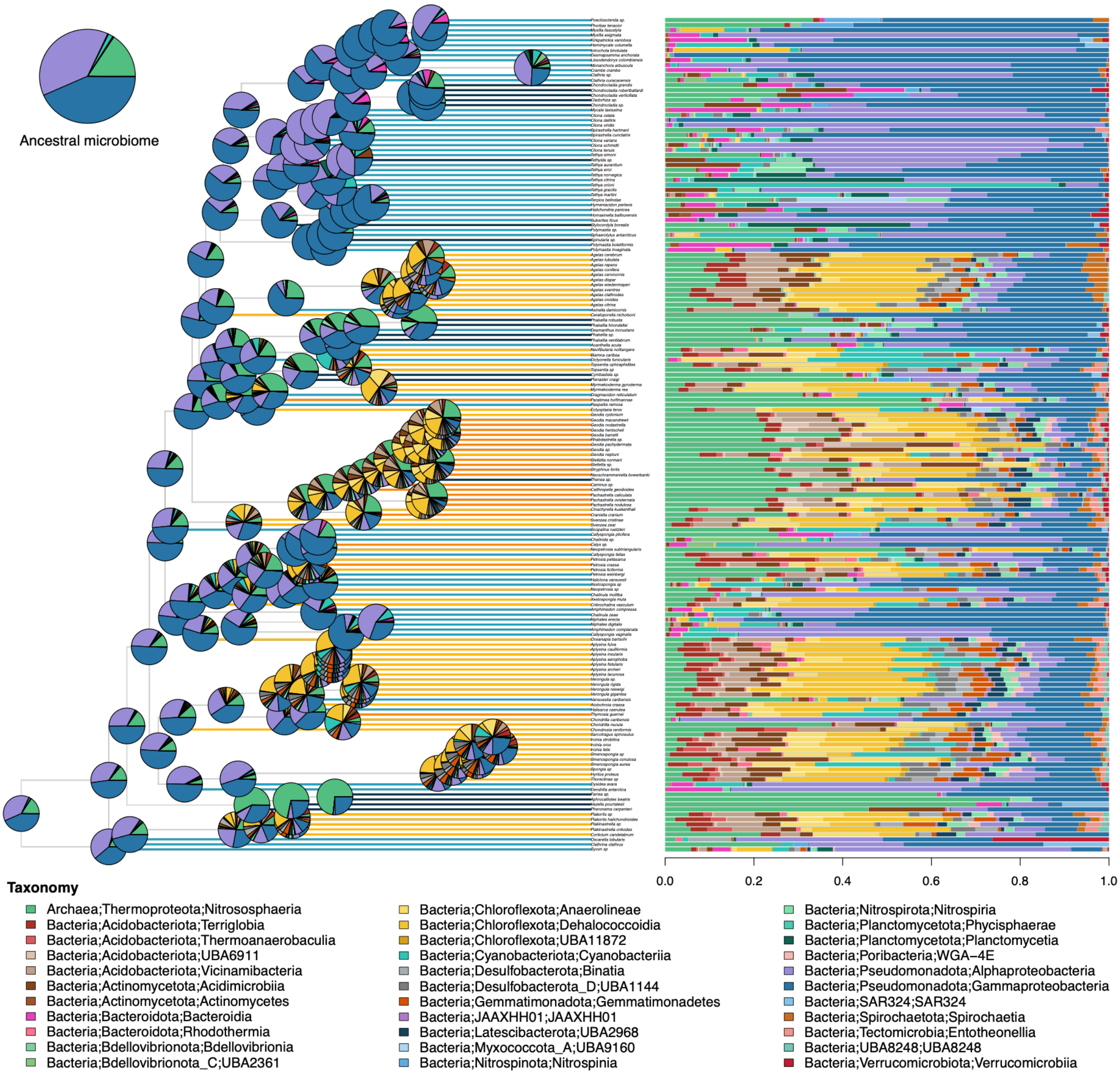


**Fig. S2. Ancestral reconstruction of sponge microbiome composition across the sponge phylogeny.** The phylogeny represents 167 sponge species, with terminal branches colored according to host HMA/LMA status and depth preference. Pie charts at internal nodes show the reconstructed ancestral microbial community composition based on the 33 most abundant microbial taxa at class level. The size of each colored sector represents the estimated relative abundance of the corresponding microbial taxon at each ancestral node. Horizontal stacked bars provide an alternative representation of the same reconstructed ancestral microbiome composition, facilitating comparison among nodes. Colors indicate microbial taxa at class level, as shown in the legend below the tree.

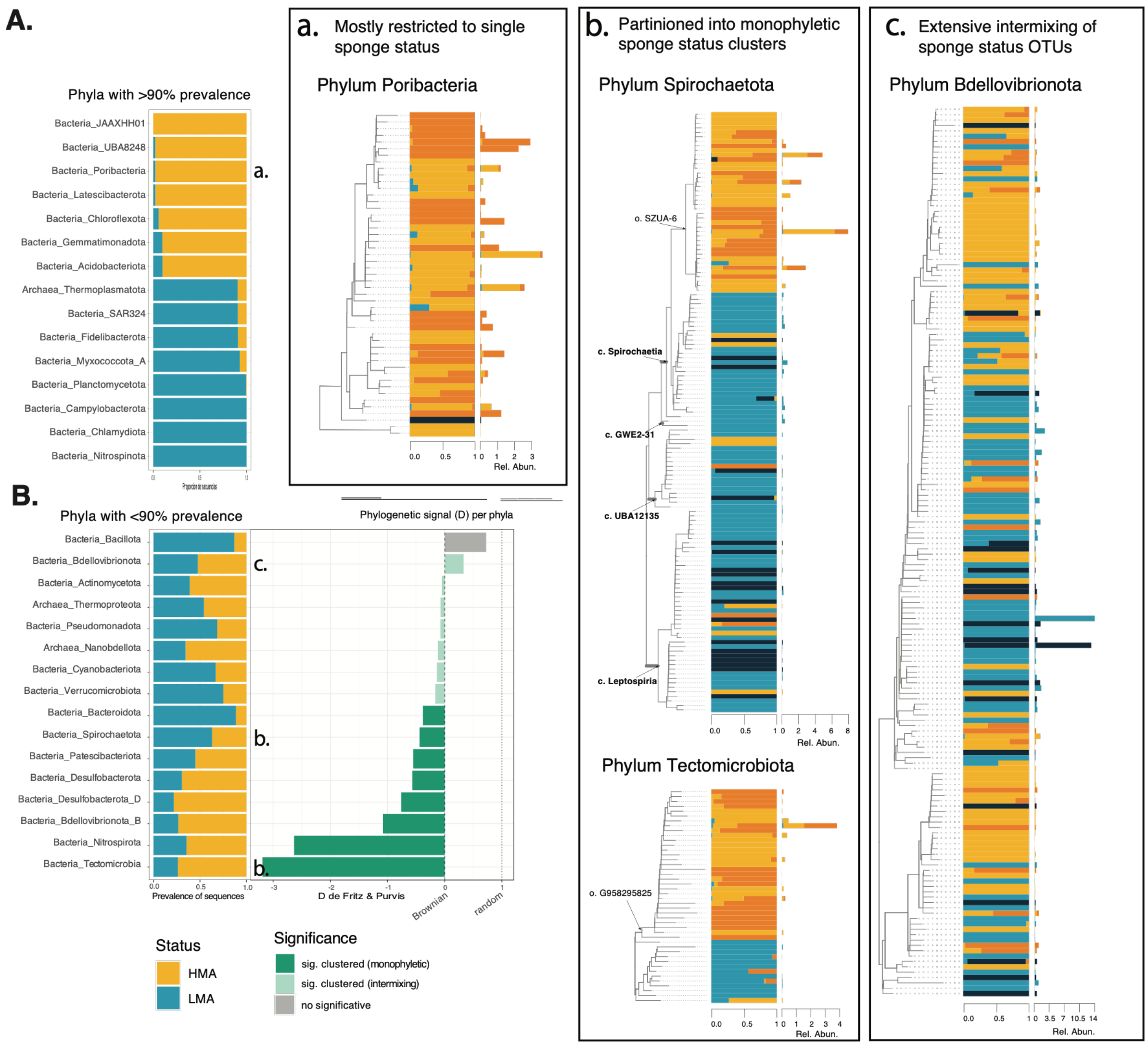


**Fig. S3. Phylogenetic organization of HMA- and LMA-associated sequences.** Phyla were classified as exclusive to one sponge status when OTU prevalence exceeded 90% (A), and as mixed when prevalence was below 90% (B). Lowercase letters in these bar plots correspond to the four phylogenetic trees shown as examples of three cases (a-c). (a) *Poribacteria* tree as an example of a phylum mostly restricted to one sponge status. (b) *Spirochaetota* and *Tectomicrobiota* trees as examples of phyla partitioned into monophyletic cluster by sponge status. (c) *Bdellovibrionota* tree as an example of a phylum with extensive intermixing of OTUs from both sponge statuses. For each branch, two bar plots are shown: the left bar plot shows the relative contribution of each OTU across the four sponge groups, while the right bar plot shows the actual relative abundance of each OTU within each group.

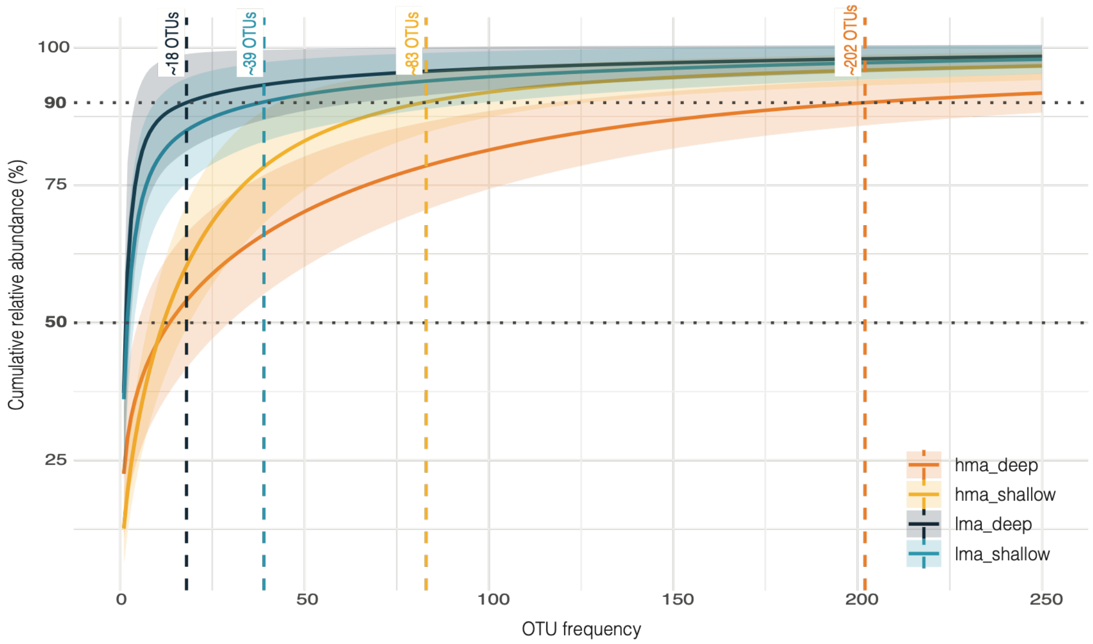


**Fig. S4. Cumulative relative abundance curves of OTUs across groups**. Most LMA species harboured one or two OTU that together accounted for 50% relative abundance, and fewer than 40 OTUs accounted for 90% of the relative abundance. In contrast, HMA species required substantially more OTUs to reach comparable abundance thresholds.

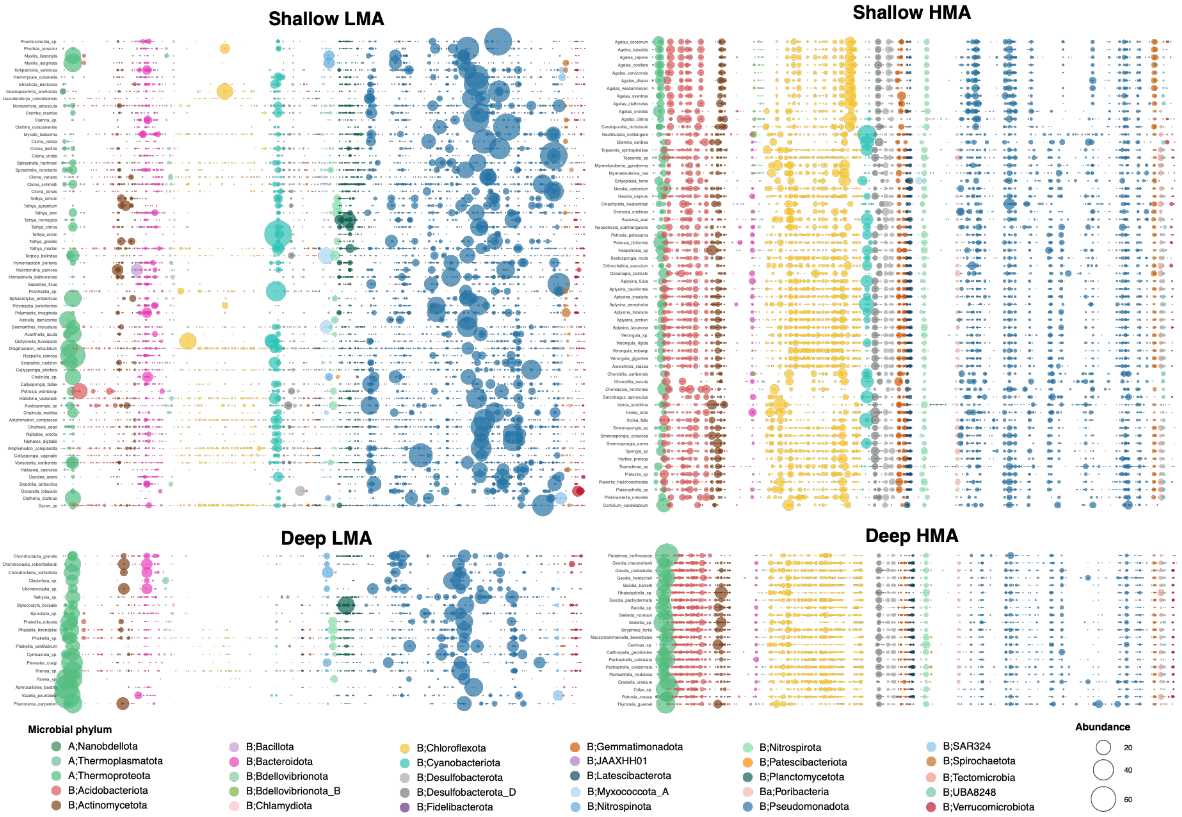


**Fig. S5. Relative abundance of individual OTUs across the four sponge groups.** Bubble plots show the relative abundance of individual OTUs, colored by phylum, for each of the four groups examined in this study. The y-axes show the sponge species included in each group, whereas the x-axes represent OTUs, which are consistent across all four plots; OTU names are omitted for clarity due to the small plot size. LMA sponges showed greater variability in OTU composition among species and strong dominance by the most abundant OTUs. In contrast, HMA sponges showed more uniform communities, with more even relative abundances and less pronounced dominance of individual OTUs.

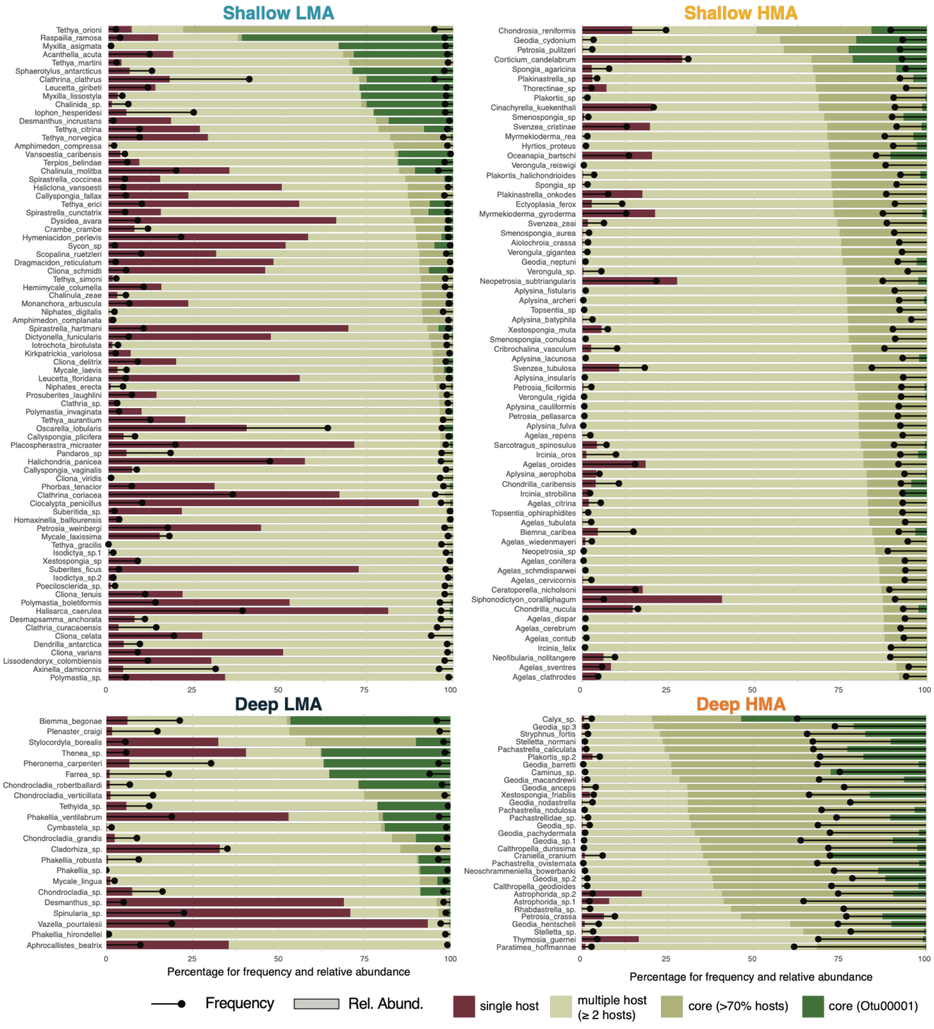


**Fig. S6. Patterns of host specificity.** Specialists (OTUs found in a single host species), shared OTUs (found in ≥ 2 host species) and core OTUs (found in >70% of host species) are shown for each sponge species within each group. The core microbiome is defined as the set of OTUs shared among >70% of host species within each group (shallow LMA, deep LMA, shallow HMA, deep HMA), whereas specialists are OTUs unique to a single species across the entire dataset. Defining specialists within each group yielded similar results. Specialist OTU frequencies are shown as lollipops extending from the left of each bar, and core OTU frequencies as lollipops extending from the right. Relative abundances are represented by bars, with OTU000001 (i.e. the most prevalent and abundant OTU) highlighted in dark green within the core microbiome.

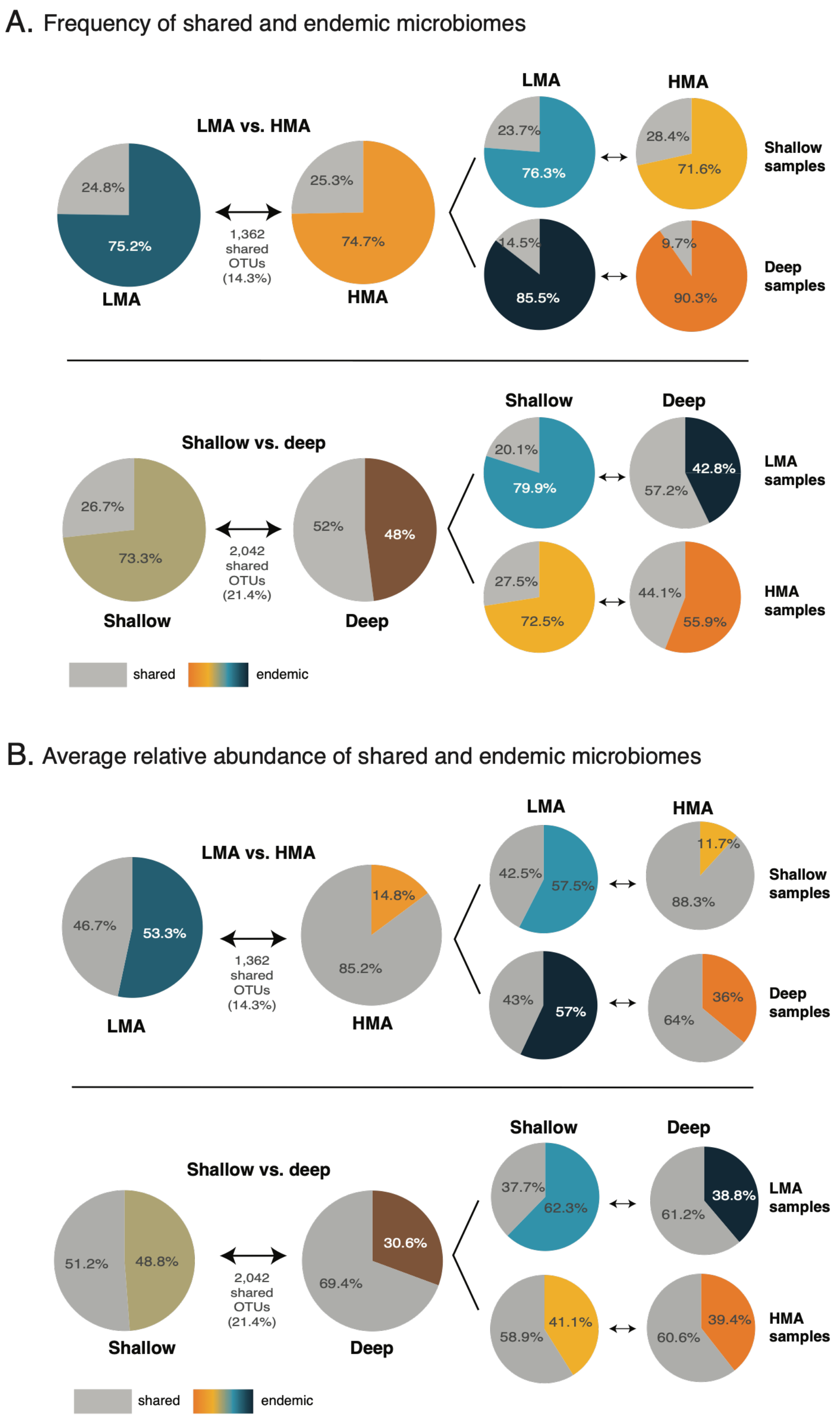


**Fig. S7. Proportions of generalist and endemic OTU fractions across sponge types and depth categories.** The upper panel compares generalist and endemic OTUs between sponge status categories, further split by depth on the right-hand side; the bottom panel compares generalist and endemic OTUs between shallow and deep environments, further split by sponge status on the right-hand side.

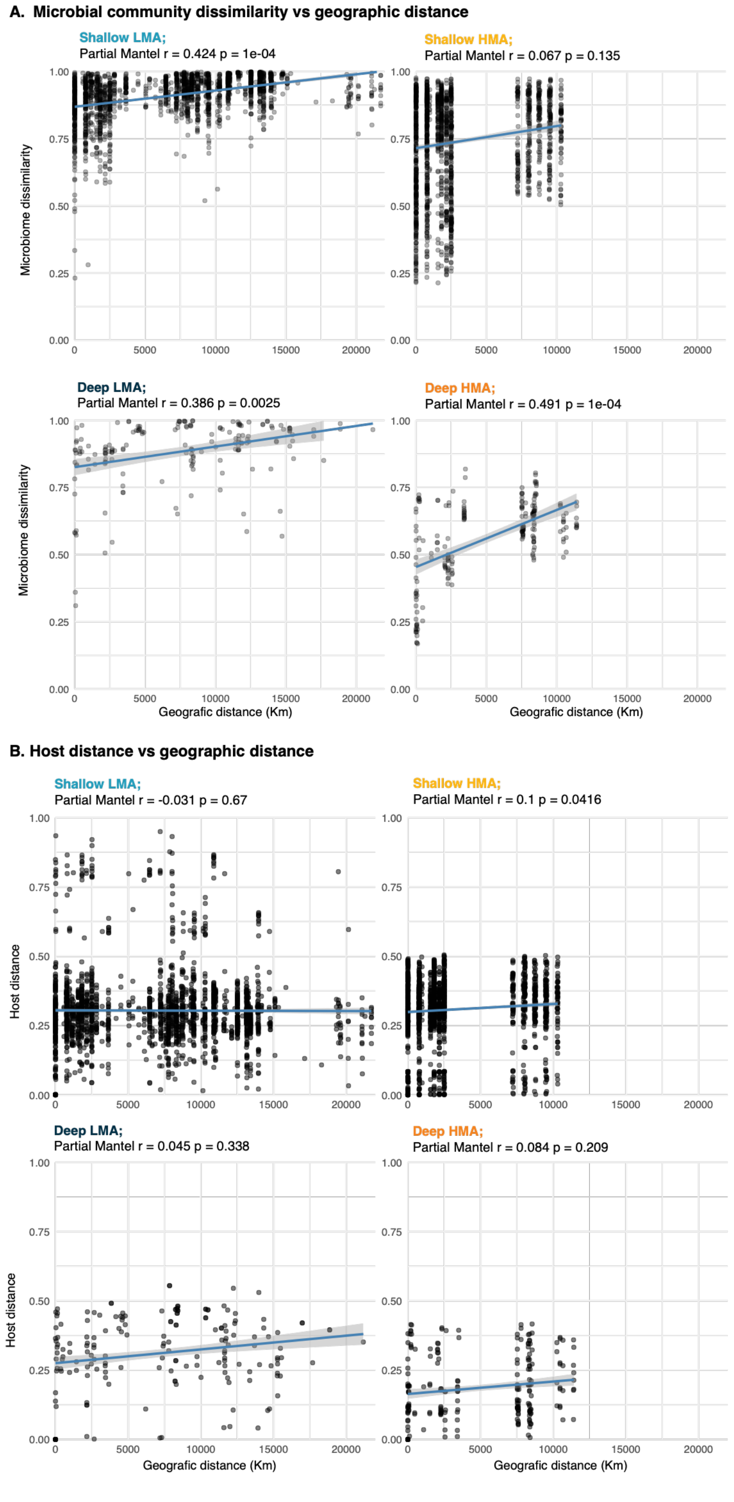


**Fig S8. Mantel tests examining the relationships among microbiome, host phylogenetic, and geographic distances. (A)** Correlation between microbiome dissimilarity and geographic distance as oceanic lest-coast distances avoiding land, across the four sample groups. **(B)** Correlation between host phylogenetic distance and geographic distance across the four sample groups. Mantel test statistics (Pearson's r) and p-values are indicated above each plot. Lines represent linear regressions with standard error (SE).

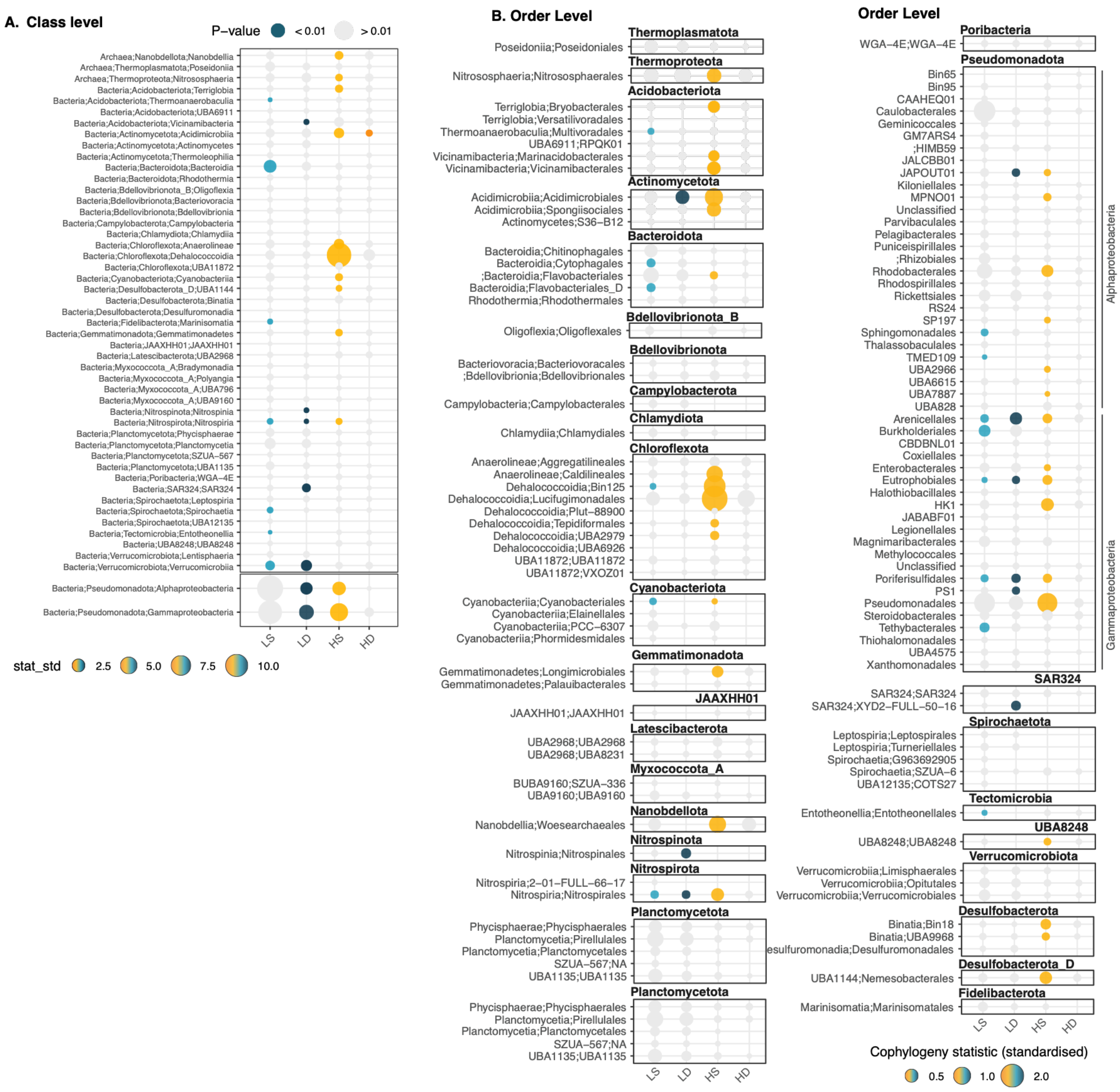


**Fig. S9. Cophylogeny analyses at additional taxonomic levels.** Cophylogeny between sponge hosts and their microbial symbionts was assessed at the class and order levels. Bubble size represents the standardized ParaFit test statistic, and colored bubbles indicate statistically significant associations (p < 0.01); non-significant associations are shown in grey.

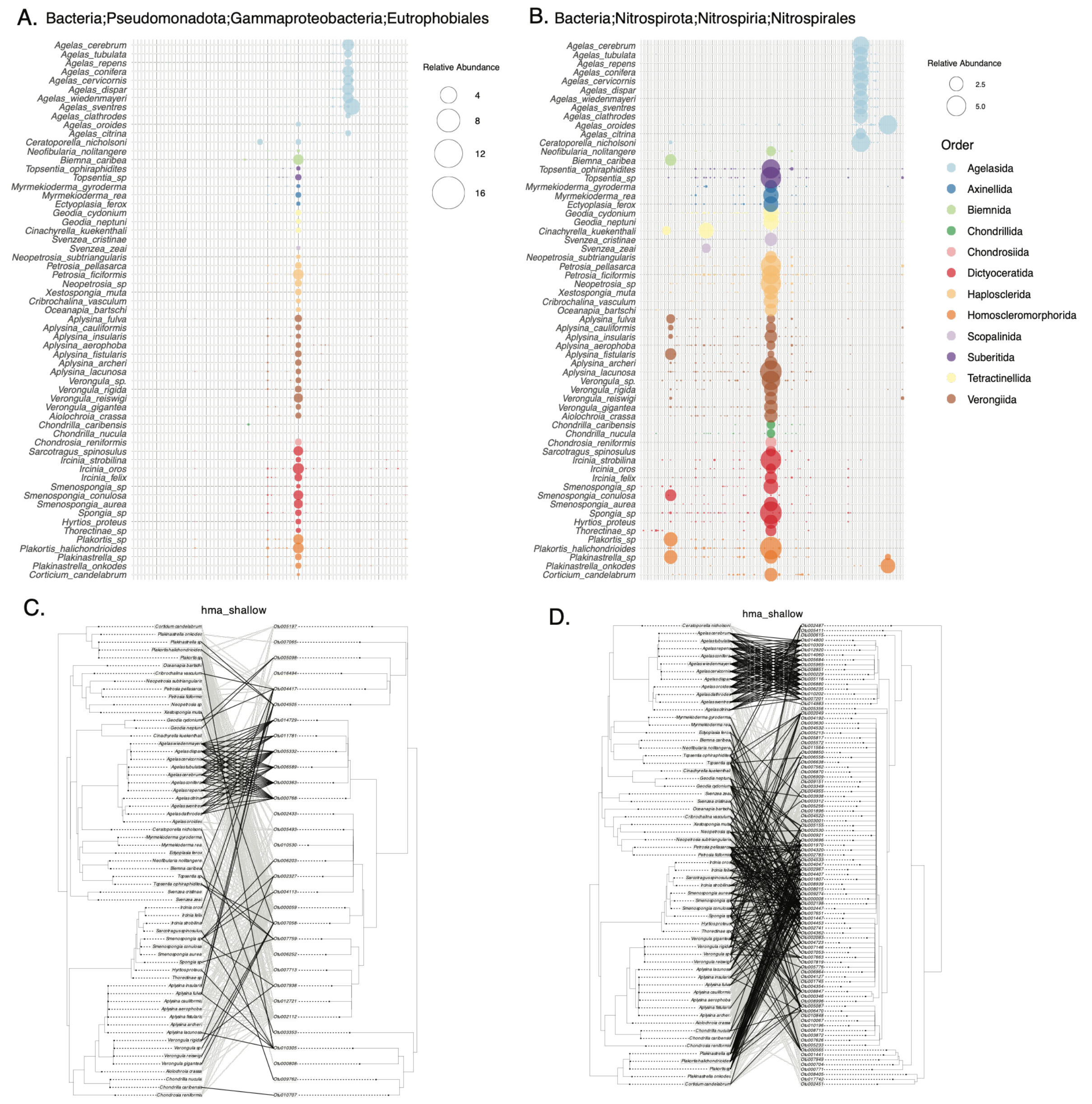


**Fig. S10. Differential microbiome composition in Agelasidae. (A–B)** Examples of specific host–microbiome associations in *Agelasidae* species, showing distinct associations with individual OTUs. **(C–D)** Phylogenetic congruence between hosts and symbionts, highlighting unique associations between *Agelasidae* hosts and phylogenetically distant symbionts.
